# Single-cell RNA sequencing identifies response of renal lymphatic endothelial cells to acute kidney injury

**DOI:** 10.1101/2023.06.09.544380

**Authors:** Heidi A. Creed, Saranya Kannan, Brittany L. Tate, Priyanka Banerjee, Brett M. Mitchell, Sanjukta Chakraborty, Joseph M. Rutkowski

**Affiliations:** Department of Medical Physiology, Texas A&M University School of Medicine, Bryan, TX 77807, USA

**Author notes:** Correspondence: Joseph M. Rutkowski; Department of Medical Physiology; Texas A&M University School of Medicine; 8447 Riverside Parkway; Bryan, TX, 77807 USA.

**Keywords:** lymphatic endothelial cells, acute kidney injury, single cell sequencing, lymphangiogenesis

## Abstract

The inflammatory response to acute kidney injury (AKI) likely dictates future renal health. Lymphatic vessels are responsible for maintaining tissue homeostasis through transport and immunomodulatory roles. Due to the relative sparsity of lymphatic endothelial cells (LECs) in the kidney, past sequencing efforts have not characterized these cells and their response to AKI. Here we characterized murine renal LEC subpopulations by single-cell RNA sequencing and investigated their changes in cisplatin AKI. We validated our findings by qPCR in LECs isolated from both cisplatin-injured and ischemia reperfusion injury, by immunofluorescence, and confirmation in *in vitro* human LECs. We have identified renal LECs and their lymphatic vascular roles that have yet to be characterized in previous studies. We report unique gene changes mapped across control and cisplatin injured conditions. Following AKI, renal LECs alter genes involved endothelial cell apoptosis and vasculogenic processes as well as immunoregulatory signaling and metabolism. Differences between injury models are also identified with renal LECs further demonstrating changed gene expression between cisplatin and ischemia reperfusion injury models, indicating the renal LEC response is both specific to where they lie in the lymphatic vasculature and the renal injury type. How LECs respond to AKI may therefore be key in regulating future kidney disease progression.

## INTRODUCTION

Acute Kidney Injury (AKI) affects up to 50% of ICU patients and is a significant contributor to the future development of chronic kidney disease (CKD)^1–3^. The initial injury extent, inflammatory resolution, and specific immune responses may play a role in the AKI-to-CKD transition^4, 5^. Increasingly, studies have demonstrated that the renal lymphatic vasculature expands in response to the increased expression of lymphangiogenic ligands upon injury and may regulate the AKI immune response through antigen, solute, fluid, and immune cell transport^6–9^. Lymphatic endothelial cells (LECs) have also demonstrated direct immunomodulatory roles during inflammation^10, 11^. Due to the relative sparsity of renal LECs compared to other cells, however, the heterogeneity and specific molecular roles of renal LECs in AKI have remained largely unexplored as the cells have been under-characterized or excluded in past sequencing efforts^12–14^. Several studies have characterized the heterogeneity of LECs along the lymphatic network from the capillary tips to conducting vessels and highlight the fluid and solute transport and immunomodulatory roles of LECs in other tissues^11, 15^.

The general anatomy and physiology of the renal lymphatic vasculature have been elegantly reviewed and characterized previously^16–18^. The network develops from the renal hilus in a vasculogenic manner similar to the dermis^16^. The adult renal lymphatic network in the mouse was recently elegantly visualized following tissue clearing and confirmed the appearance of valved collecting lymphatic vessels along the renal arteriole network and limited lymphatic capillaries throughout the renal parenchyma^19^. The molecular and genetic heterogeneity of renal LECs, however, is largely unknown.

Following an AKI, lymphangiogenesis is initiated in response to inflammatory stimuli and the upregulation of lymphatic growth factors. Previous studies investigating whether renal lymphangiogenesis is beneficial to the recovery of renal function and clearance of immune cells has remained inconclusive^20^. Some studies have indicated renal lymphangiogenesis as being a potential driver of kidney inflammation^21^. However, other evidence suggests lymphangiogenesis has a largely beneficial effect on the recovery of kidney function^6, 9, 22^. For example, our past work initiating renal lymphangiogenesis prior to injury altered the CD4:CD8 T cell ratio and demonstrated improved long-term renal functional and fibrotic outcomes^22^. These findings indicate that renal lymphatics and LECs may respond dynamically to various injury stimuli and directly alter the post-injury immune response through lymphangiogenesis and changes in LEC biology.

In this study, we isolated murine renal LECs and utilized scRNA sequencing to identify LEC vessel subpopulations in quiescence and determined distinct molecular responses for each subset in response to cisplatin-induced AKI. We analyzed global renal LEC specific roles, evaluated their molecular identities and functions, and assessed how renal LEC functions alter gene and pathway expression following an AKI. Renal LEC composition was identified to have few lymphatic capillaries and the network was strongly lymphangiogenic in response to AKI. The predominant immunomodulatory pathways in AKI were found to be in LEC-T cell interactions. These findings thus provide a novel characterization of the renal lymphatic vasculature and identify potentially how LECs impact the progression of AKI.

## METHODS

### Animal Injury Models

All experiments were performed in male C57BL6/J mice aged 10-12 weeks of age and assignment to injury was not randomized. For cisplatin AKI, a single intraperitoneal injection of cisplatin (Vet Tech, 68001-283-27) or saline control was administered to mice via a single i.p. injection at a dose of 20mg/kg (n= 5 mice per group). Bilateral ischemia reperfusion injury procedures were conducted as previously described by Texas A&M University Comparative Medicine Program veterinary staff^22^. Briefly, mice were anesthetized by ketamine/xylazine intraperitoneal injection and both renal pedicels were revealed, and blood flow halted by atraumatic vascular clamps (Fine Science Tools, 18055-05) for 20 minutes. Mice were maintained at 37°C throughout the procedure and renal ischemia was confirmed visually through loss of kidney color. Restoration of kidney color confirmed renal reperfusion prior to suturing. All mice were monitored daily to assess body weight loss, behavior, or major signs of distress. Mice were euthanized 72-hours following AKI (n=5 mice per group). Mice were perfused via the left ventricle with 5 mL of ice cold HBSS (14025092, Gibco) after which perfused kidneys were collected and placed on ice in HBSS. To prevent batch effects all kidneys were collected and processed on the same day. Mice were provided *ad libitum* access to water and standard chow throughout the study. All animal protocols and housing for the study were approved by the Institutional Animal Care and Use Committee at Texas A&M University.

### Cell Isolation and Tissue Collection

For each condition, five mice were utilized in single cell enrichment and sequencing experiments as determined by previous isolation and flow cytometry experiments. Kidney capsules were retained, and kidneys were minced into small pieces and placed into 5mL digestion buffer (15mg/mL Collagenase D (Roche Applied Science, 11088858001), 10mg/mL Collagenase Type 4 (Gibco, 17-105-019), 2 mg/mL Dispase II (Roche Applied Biosciences, 4942078001), BSA Fraction V (Fisher Scientific, BP1605-100), and HBSS (ThermoFisher Scientific, 14025092). The samples were then incubated in a 37°C water bath with manual agitation every 10 minutes. Digestion was halted by adding 10mL of ice cold HBSS. Digested samples were filtered through both 70µm and 40 µm filters. Endothelial cells were enriched magnetically using a mouse CD31 biotinylated antibody (BD Pharmingen, 553371) and a cleavable Biotin Binder Kit (Invitrogen, 11533D). After the linker was cleaved, the isolated cells were subsequently enriched using an anti-mouse podoplanin (Pdpn) biotinylated antibody (BAF3244, R&D Systems) as above. CD31+Pdpn+ cells were placed into ice cold RPMI 1640 + 10% FBS prior to FACS confirmation of CD45-CD31+Pdpn+ cells. Cell fractions were resuspended in in RPMI1640 +10% FBS, counted, and diluted to a concentration of ∼1,000 viable cells/μL.

### scRNA sequencing Bioinformatics Pipeline and Analysis

#### Library Generation and Quality Control

CD31+Pdpn+ cell count and viability were confirmed on the ThermoFisher Countess and the Moxi Go II (Orflo, MXG102). Samples were then loaded onto the Chromium X Controller using the Chromium Next GEM Single Cell 3’ Reagent Kit V3.1. Sample cDNA quality was checked using the Agilent High Sensitivity D5000 ScreenTape assay on the Agilent 4200 TapeStation System (Agilent Technologies, G2991BA). cDNA samples were then quantified and normalized prior to the library sequencing. Libraries were sequenced on a NovaSeq S4 Flow cell according to 10x Genomics recommendations. Single-cell RNA sequencing reads were processed using 10x Genomics cloud analysis. Briefly, fastq files were uploaded to the 10x Genomics Analysis cloud and aligned to the reference mouse transcriptome (mm10) using Cell Ranger Count v6.0.1. a digital gene expression matrix, including raw unique molecular identifier (UMI) counts were assigned to each cell in the respective sample.

#### Bioinformatics data processing and in Silico selection

Gene expression matrices generated using the CellRanger software (10x Genomics) and raw data reads were processed in R (version 4.2.1). Cells then underwent quality control steps prior to downstream analyses. Cells with fewer than 200 detected genes, more than 4,000 genes, or more than 60% mitochondria reads were excluded. Count data was normalized using the NormalizedData function in the Seurat package (version 4.0) and highly variable genes were identified with FindVariableFeatures function. Comparable populations were identified in scRNA sequencing and anchored in both control and injury conditions. The ScaleData function was utilized to scale expression of genes to provide equivalent weight to all genes. Data was then summarized using principal component analysis (PCA) in the Seurat package followed by visualization with Uniform Manifold Projection (UMAP). Cells were clustered according to their gene expression profile using the FindClusters function in Seurat with a Canonical Correlation Analysis (CCA) clustering resolution of 1. CCA resolution of 1 was set after evaluation of PCA elbow plots, UMAP assessments of cell quality effects, and presence of distinct top gene markers present for each population. Clusters were identified by expression of known vascular endothelial cell (EC) marker genes *Emcn, Cd34,* and *Kdr,* hybrid vascular markers *Emcn, Flt4,* and lack of *Pdpn,* lymphatic marker genes *Pdpn, Prox1, Flt4,* and *Sox18,* and immune cell gene *Ptprc*. After identification of cluster identities and initial global analysis all non-lymphatic cell clusters were removed from further downstream analyses. Gene Ontology (GO) and MCODE analyses were processed and visualized using the Metascape tool^23^.

#### Subcluster Analysis and Identity Assignment

Control and cisplatin injured LEC clusters were subclustered using Seurat’s subset function. A new Seurat object was generated and processed as described above in bioinformatics data processing. Four new clusters were identified using the FindClusters function with a CCA resolution of 0.25. New LEC subcluster differentially expressed genes were determined using the FindAllMarkers function with an average log fold change of ≥0.2. New LEC subclusters identities were assigned using previously published lymphatic data sets genes which correspond to vessel regional identity^15, 24–26^. For injury comparisons, a LEC subset was generated as described above using the subset function. Control and cisplatin injury LEC subset genes were compared using the FindMarkers function and top significantly upregulated and downregulated between conditions were identified. GO analyses were generated as described above. Violin plots of significantly altered immune genes were generated using the VlnPlot function in the Seurat package.

### In Vitro Human Dermal Lymphatic Endothelial Cell (HDLEC) Cisplatin Treatment

Human dermal lymphatic endothelial cells (HDLECS) were purchased from Promocell (C-122217) and maintained in complete endothelial basal media (C-22022, Promocell) at 37°C, 5% CO2 conditions as previously described^27^. Confluent HDLECs were then treated with either a saline control, 4 µg/mL, or 8 µg/mL cisplatin for 24 hours according to previously described studies^28–30^. A dose of 12 µg/mL cisplatin was lethal to HDLECs. Cells were then isolated at the end of the 24 hours treatment period and total RNA was extracted using the PureLink™RNA Mini Kit (12183018A, Invitrogen).

### In vivo LEC RNA isolation for validation

LEC enrichment was performed as above and the RNA from the LEC-enriched endothelial cell isolation was isolated from mouse kidneys of control, cisplatin AKI, and IRI conditions using the Direct-zol RNA Miniprep Plus kit (R2072, Zymo) following the manufacturer’s instruction.

### qPCR

Complementary DNA was synthesized from 0.5μg total RNA for all *in vivo* and *in vitro* experiments using the iScript cDNA kit (1708891, Bio-Rad). Five µL qPCR reactions were carried out using iTaq Universal SYBR Green Supermix (1725124, Bio-Rad) Reactions were run in duplicate on a QuantStudio6 Flex Real-Time PCR system (Applied Biosystems). Fold changes compared to control samples were calculated using the 2 ^ΔΔCT^ method with *Ubc* and *ACTB* as endogenous control for the *in vivo* and *in vitro* samples respectively (Supplemental Table 1). All *in vivo* data from the CD31+Pdpn+ cells were normalized by a lymphatic endothelial cell specific marker *Flt4* (VEGFR-3), 2 ^ΔΔCT^ values of all the genes were divided by the 2 ^ΔΔCT^ value of *Flt4*.

### Immunofluorescence

For tissue immunofluorescence analyses kidneys were formalin fixed for 24 hours and, following a sucrose gradient, embedded in OCT (T4583, Tissue-Tek) for sectioning. 10µM kidney sections were incubated overnight at 4°C with selected primary antibodies (Supplemental Table 2). Sections were then washed with PBS and then blocked in 10% donkey serum for 30 minutes. Sections were then incubated with secondary antibodies. Slides were mounted with Prolong Gold Antifade with DAPI (OB010020, SouthernBiotech) and visualized with an Olympus BX51 fluorescent microscope with an Olympus Q5 camera. Representative images were captured at figure indicated magnification.

### Statistical Analyses

Statistical analysis of qPCR data was performed using GraphPad Prism software (version 9.5.1). An unpaired *t*-test with unequal variance (Welch) was utilized and *p* value <0.05 was considered significant.

### Data and Code Availability

The single-cell RNA sequencing data generated in this study is deposited in GEO (https://www.ncbi.nlm.nih.gov/geo/) with accession number: *(on publication)*. No unique code was generated from this study.

## RESULTS

### Renal LEC isolation and scRNA sequencing identification of LEC populations

Lymphatic endothelial cells are few in number compared to other cells in the kidney and often missing in past scRNA sequencing studies^13, 14^. To identify the heterogeneity of LECs in the kidney, we collected both kidneys from 5 mice, enriched LECs, and performed scRNA sequencing (Figure 1A). Enrichment of renal LEC identity was confirmed prior to scRNA sequencing library preparation by flow cytometry. Based on the isolation protocol utilized, approximately 99% of cells were CD45-. Of these, approximately 60% highly expressed CD31 and Pdpn with nearly all of these cells also demonstrating LYVE-1 on their surface (Figure 1B). Sequenced samples were thus approximately 60% LECs by classically described surface expression. Unsupervised clustering analysis of the total cell pool assigned seven unique cell clusters to the anchored data (Figure 1C). After quality control filtering and exclusion of contaminating cell types, we obtained a total of 2,323 cells for analysis. To discriminate LEC populations from other CD31+PDPN+ cells sequenced, gene expression was limited to cells expressing the following markers: vascular genes *Kdr, Emcn,* and *Cd34* and LEC genes *Prox1, Flt4,* and *Sox18* (Figure 1D, Supplemental Figure 1). As expected, scRNA sequencing populations were a mix of both LECs and other vascular phenotype-like cells. It is noted that previous studies have indicated the presence of other Pdpn+ expressing cells in the medullary region, such as urothelial cells and ascending and descending loop epithelium, in the kidney^17, 31–33^. Three LEC populations were thereby identified in addition to three vascular, and one immune population. LEC1 and LEC2 genetic populations were quite similar, while LEC3 was genetically more like the “hybrid endothelium” attributed populations.

**Figure 1:**
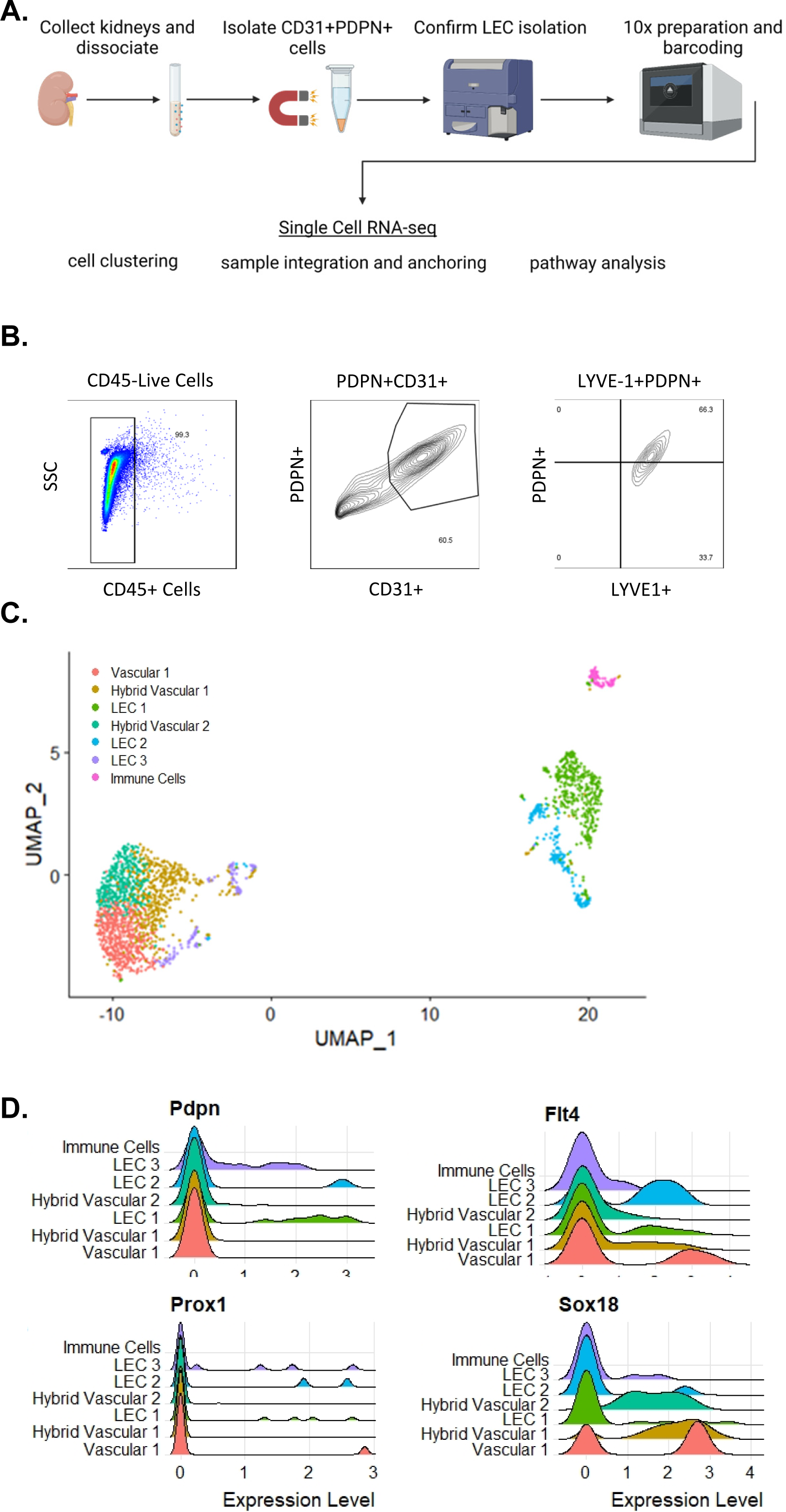
Isolation and Identification of renal lymphatic endothelial cells. A) Diagrammatic outline of lymphatic endothelial cell (LEC) isolation method and follow on bioinformatics analysis. B) Flow cytometric verification of renal lymphatic endothelial cell populations captured with magnetic enrichment as verified by gating of CD45-live cells, PDPN+CD31+, and LYVE1+PDPN+ C) Anchored clustering analysis of renal LEC single cell data sets identifies seven cell populations through canonical correlation analysis (CCA) unsupervised clustering. D) Lymphatic identity of cell populations verified through expression of lymphatic genes.

### Renal LEC heterogeneity is similar to lymphatic structural components from other tissues

To define each cluster globally, we assessed the top 5 differentially expressed genes that identified each unique cluster in the full cell population (Figure 2A, B, Supplemental Table 3). We identified that LEC1 and LEC2 populations were distinct in quiescent conditions and tested if these populations could be assigned to established lymphatic vessel components based on their differentially expressed genes. LEC1 was characterized by heightened expression of genes *Fabp4, Plvap, Rsad2,* and *Kdr,* while LEC2 was defined by *Eln, Plxnb2,* and *Fbln5* (Figure 2A). Genetic expression of *Fabp4*^24, 34^*, Plvap*^34^, and *Kdr*^35^ have been previously well-annotated in lymphatic vessels. LEC2 genes *Flbn5* and *Eln* have been described in cases of lymphatic hypoxia^36^ and *Plxnb2* is a receptor for SEMA4C on LECs^37^. In differentiating these LEC populations, there was no discernible association to specific lymphatic vessel anatomical features (e.g., lymphatic valves) or developmental lineage (e.g., sprouting versus lymphangioblasts) from global analysis. LEC3 cells in quiescent conditions were sufficiently different and small in population number such that they were not analyzed further as LECs or compared in injury conditions (Supplemental Table 3).

**Figure 2:**
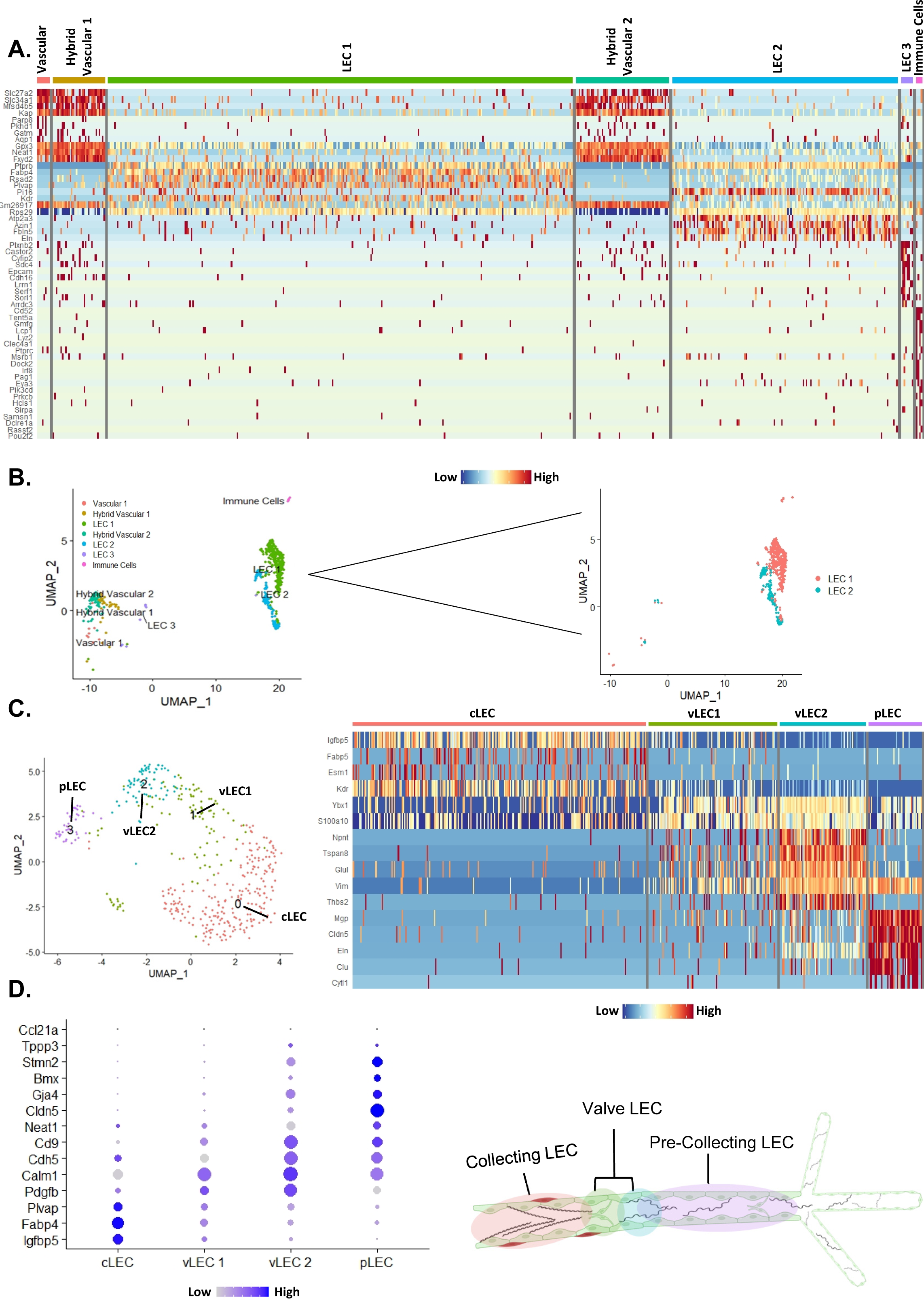
Genetic signatures of renal lymphatic endothelial cells. A) In control conditions lymphatic endothelial cell (LEC) populations 1 and 2 comprised a bulk of the population and isolated as a subset for LEC specific analyses. In control conditions 458 total LECs were utilized in downstream analyses. Scaled expression level heatmap of top 5 genes from the identified cell clusters. B) UMAP of control isolated cell populations and selected LEC subset (composed of LEC 1 and LEC2) utilized for subsequent downstream analysis. C) Expression level scaled heatmap of top 5 marker genes in LEC subcluster phenotypes. LEC subclusters differentiate into four distinct anatomical locations. D) Previously published gene markers used to identify distinct lymphatic vessel subpopulation LECs. E) Diagrammatic depiction of the anatomical location of captured LEC populations.

To determine if the LECs were associated with a lymphatic vessel structural feature and assign their identities as such, we performed subset analysis on LEC1 and LEC2 (Figure 2B). LEC subset analysis revealed four subclusters and new defining differentially expressed genes (Figure 2C, Supplemental Table 4). We utilized previously published lymphatic markers to identify the following features: collecting LECs (cLECs) *Igfbp5*^38^, *Fabp4*^24^, and *Plvap*^34^, valve LECs (vLECs) *Calm1*^38^, *Pdgfb*^24^*, and Neat1*^38^, and pre-collecting LECs (pLECs) *Cldn5*^24^*, Gja4*^24^, and *Bmx*^24, 39^. cLECs expressed genes as defined above and vLECs split into two distinct populations. vLEC1 was defined by listed vLEC markers while vLEC2 shared expression of vLEC markers in addition to expression of pLEC and capillary LEC markers *Stmn2, Tppp3,* and low expression of *Ccl21a*^24, 40^ (Figure 2D, E). Interestingly, we did not isolate or identify a distinct lymphatic capillary population in our data set, however, past studies examining murine lymphatic anatomy have found sparse lymphatic capillary structures within the kidney ^16, 19, 41, 42^.

### Renal LEC gene expression changes in response to cisplatin AKI

In AKI, whether the lymphatic system and LECs hold a beneficial or detrimental role in resolution or progression following injury is not yet clear. To characterize renal LEC transcriptional adaptation to AKI, mice were injected with a single high dose of cisplatin and renal LECs were isolated 72 hours post injury. After injury there was an increase of total number of Pdpn+ expressing cells in the kidney. This was not due to lymphatic expansion (appreciable lymphangiogenesis is only significant around 7 days post cisplatin injury^9^). The populations of LEC1 and LEC2 were reduced in numbers compared to control conditions (Figure 3A). LECs were not lost; rather, the number of “hybrid” cells increased relative to the total. De novo Pdpn expression was found in the inner medulla (Supplemental Figure 2A). Pdpn expression upon injury has been previously reported in medullary collecting duct cells: Pdpn expression in the inner medulla was negative for potential vasa recta or endothelial markers PLVAP and SLC14a1 by immunofluorescence confirming their lack of “LEC-ness” (Supplemental Figure 2B, C). An increase in isolated immune cells was also noted in AKI (Figure 3B).

**Figure 3:**
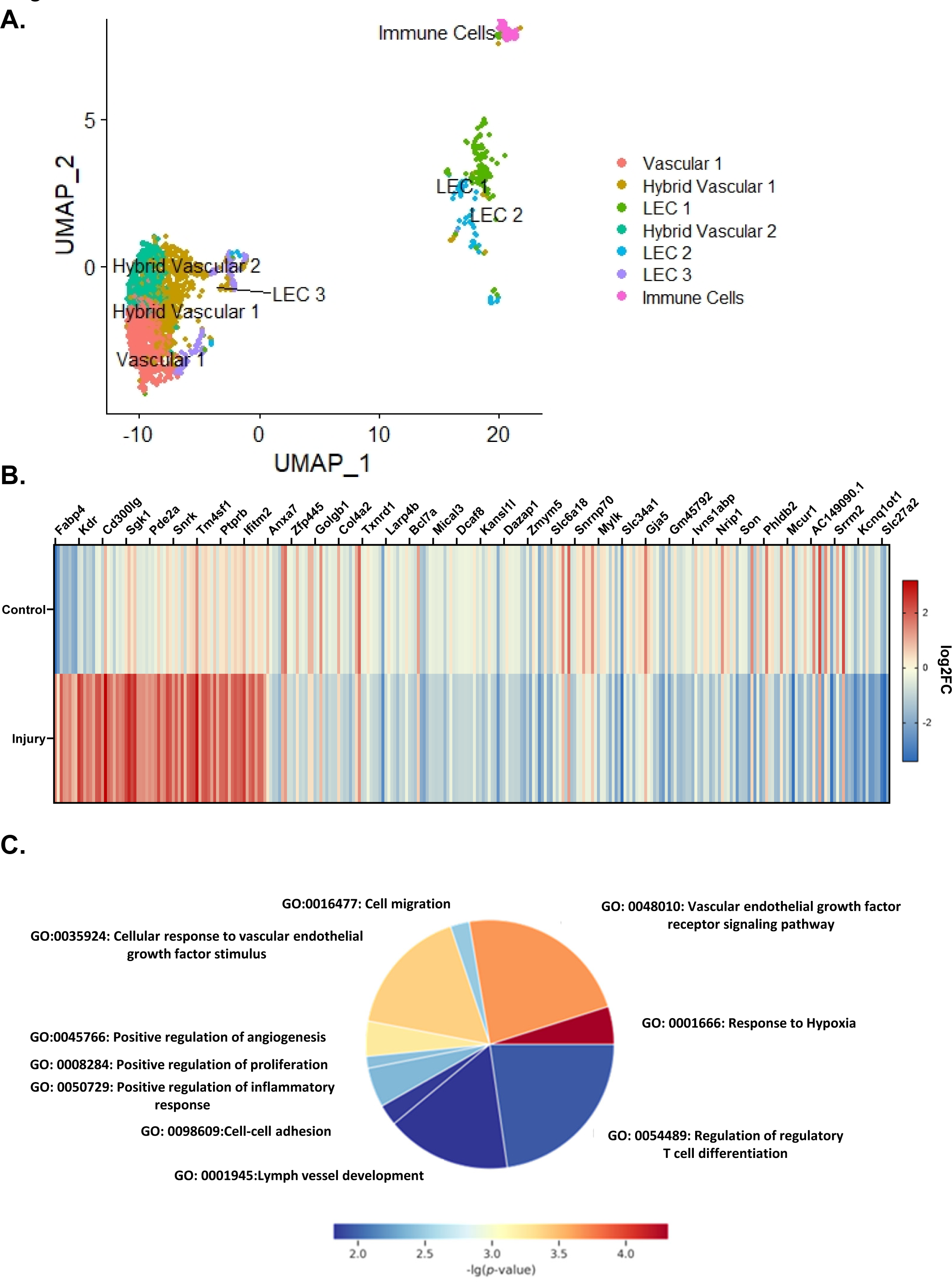
LEC injury response demonstrates altered molecular roles. UMAP of Cisplatin injury conditions demonstrated a shift towards increased podoplanin expressing vessels and a distinct shift in LEC 1 and LEC 2 populations. A) Heatmap of 283 most altered LEC subset genes between control and injury conditions. Fold change indicates increased (red) or reduced (blue) expression for indicated gene. Adjusted p< 0.05. B) Pie chart of top 25 upregulated injury genes when compared to control conditions indicate indicates top changed genes are associated with biological processes such as: increased angiogenic signaling, lymphatic vessel development, and increased inflammation related processes.

To understand how LECs respond to AKI we compared LEC subset gene expression differences between injured and quiescent conditions (Supplemental Table 5). Changes in LEC 1 and LEC 2 log_2_FC were examined briefly prior to LEC subset comparison between quiescent and injury conditions (Supplemental Figure 3 A-C). GO analyses identified the top changed injury genes when compared to quiescent conditions as *Fabp4* (log_2_FC 4.89)*, Kdr* (log_2_FC 3.24), and *Plpp3* (log_2_FC 3.19) (Figure 3C). These genes are highly involved in fatty acid transport, VEGFR signaling, and immunoregulatory interactions, roles identified in the injury GO analyses (Supplemental Figure 4 A,C). Downregulated processes included oxidative stress and redox pathways and dicarboxylic acid transport, key pathways in metabolism which are altered with cisplatin injury^43, 44^ (Supplemental Figure 4C). Genes *Cd300lg* (log_2_FC 2.37), a gene encoding an immunoglobulin like protein which plays a role in lymphocyte homing and adhesion to endothelial cells^45^, and *Tspo* (log_2_FC 2.11), a gene highly involved in immune cell processes and previously implicated in apoptosis prevention in renal ischemia reperfusion injury models^46, 47^.

To understand how highly changed genes corresponded to specific biological processes, the top 30 upregulated genes in injury conditions, when compared to quiescence, were grouped and characterized according to GO biological process. The majority of genes were associated with VEGF receptor signaling pathways, VEGF stimulus, and lymphatic vessel development: all priming LECs undergoing lymphangiogenesis. However, roughly a quarter of the genes were associated with T cell differentiation (*Ctla2a, Id1, Cd38, Rsad2)* and three additional genes were implicated in the regulation of the inflammatory response (*Plpp3, Ifitm3, Bst2)*(Figure 3D). LECs have previously demonstrated T cell related immunoregulatory roles in other disease models such as cancer, arthritis, and cardiac injury^48–51^. This data lends additional support to the finding renal LECs are undergoing proliferation and upregulating their interaction with immune cells and alter their molecular repertoire upon kidney injury.

### Renal LEC adaption to injury stimulus and anatomical composition

To assess LEC involvement with the immunological transition between innate and adaptive immune processes, genes related to innate immune response processes were selected and plotted to compare expression between control and injury conditions. Genes related to integrin signaling and the innate immune response such as *Ltbp4* and *Kitl* were decreased in LECs following injury along with *Plscr2*, a mast cell activating gene, and *Foxp1,* a negative regulator of T follicular helper cell differentiation. However, genes *Spp1* and *Igfbp7* were present almost exclusively in injury. The transcript of *Spp1*, osteopontin, acts as a key renal cytokine while the transcript of *Igfbp7*, insulin-like growth factor-binding protein 7, has been demonstrated to have roles in stimulating cell adhesion, blocking angiogenic signaling, and changes cell sensitivity to chemotherapeutic drugs^52, 53^ (Figure 4A).

**Figure 4:**
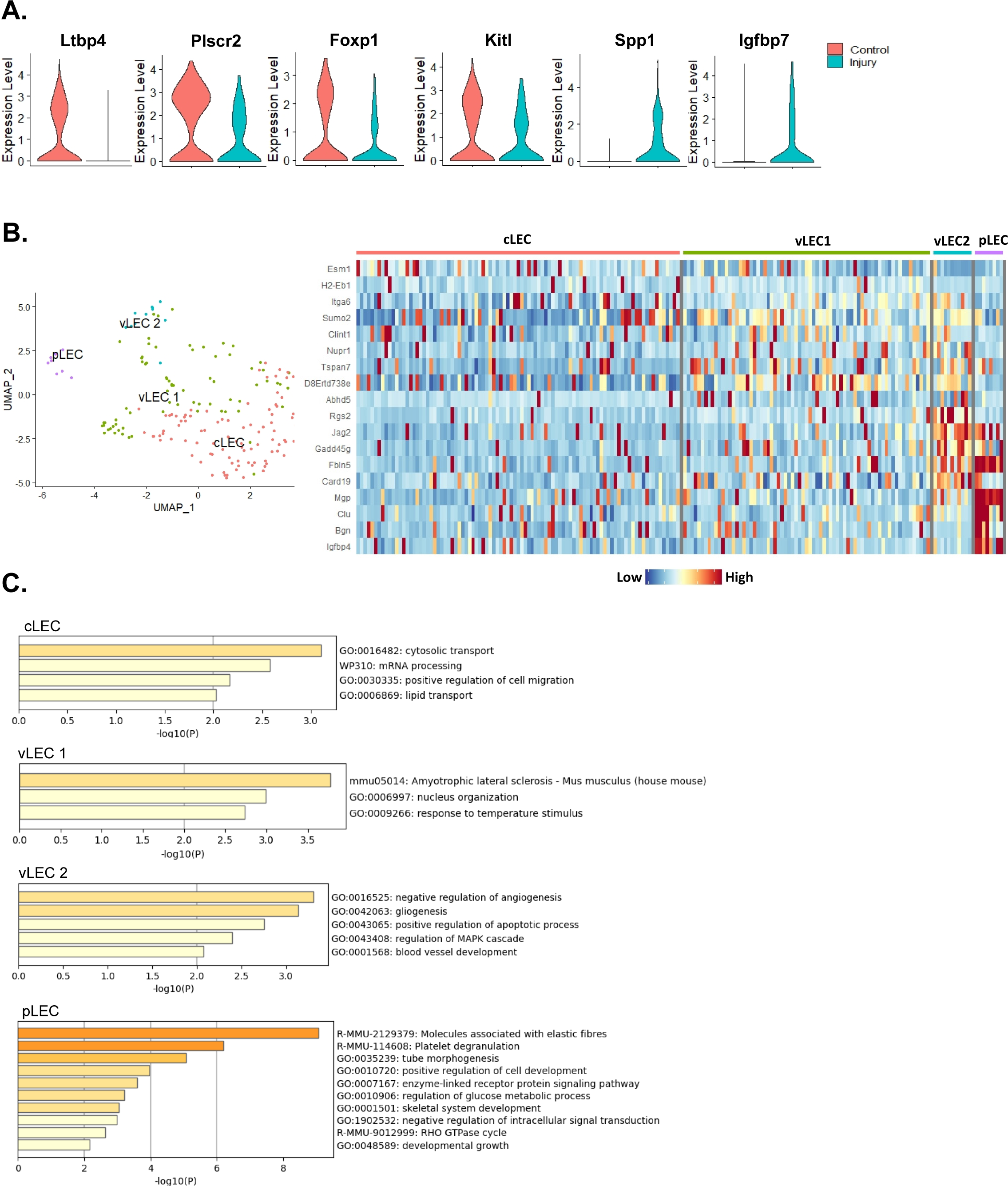
Renal LEC adaptation to injury stimulus and anatomical composition. A) Violin plots of selected immune process related genes altered between control and injury conditions. Genes selected represent changed genes between conditions which are additional related to innate and adaptive signaling processes. B) UMAP of injured LEC subset demonstrating similar anatomical populations to control conditions. Expression level scaled heatmap of top 5 marker genes in LEC subcluster phenotypes; LECs differentiate into four distinct anatomical locations. C) In injury conditions lymphatic vessel anatomical identity GO analyses identify changes in defined genetic roles.

Among the anatomical LEC subpopulations, cLEC and vLEC1 subpopulations lose the genetic resolution observed in control conditions following injury (Figure 4B). Notably, pLEC retains some defining genes from the control conditions (*Mgp, Clu*) unlike the cLEC, vLEC1, and vLEC2 populations. GO analysis was performed on LEC subpopulations to determine how injury alters the molecular functions of renal LEC anatomical locations. By GO analysis, cLECs are involved in cell migration and transport processes and vLEC populations are involved in nuclear organization, apoptotic processes, and angiogenic signaling. In injury, pLECs are more highly involved with developmental processes and metabolism (Figure 4C). Overall, this data indicates LECs maintain similar anatomical composition to quiescent LECs but alter their vasculogenic signaling and metabolism.

### Validation of renal LEC gene response to kidney injury

To validate LEC specific gene changes in injury we utilized immunofluorescent labeling and qPCR on renal LECs. We identified *Spp1*, one of the highly variable genes in injury conditions and associated with the LEC injury response to co-localize with Pdpn+ lymphatic vessels exclusively in injury conditions (Figure 5A). We did not detect immunofluorescent co-localization of Spp1 in uninjured kidneys. We further confirmed co-localization of a second gene target in injured kidneys, *Fabp4*, which was one of the most changed in the injury LEC subset (Figure 5B). We did not note co-localization of either Spp1 or Fabp4 with Pdpn+ glomeruli, indicating their Pdpn+ co-localized upregulation was lymphatic specific. As immunofluorescence options were limited, fresh LEC-enriched cell isolations were tested by qPCR in cisplatin and IRI injured LECs. Tested genes were selected based on both significance in injury and the largest log_2_FC between conditions. Gene expression between conditions was compared in a heatmap to determine the most altered genes between injury models (Figure 5C). Genes *Ptprb, Mgp, Gng11, and Vim* demonstrated a significant increase in relative expression validating scRNA sequencing findings (Figure 5D). Additionally, while not significant *Kdr* demonstrated a trend toward increased gene expression in injury by qPCR in the LEC-enriched cells. In IRI conditions, *Ptprb* was also significantly increased but *Vim* and *Fabp4* were significantly decreased indicating an injury specific effect. This data indicates that the scRNA sequencing findings are reproducible and detectible in LECs.

**Figure 5:**
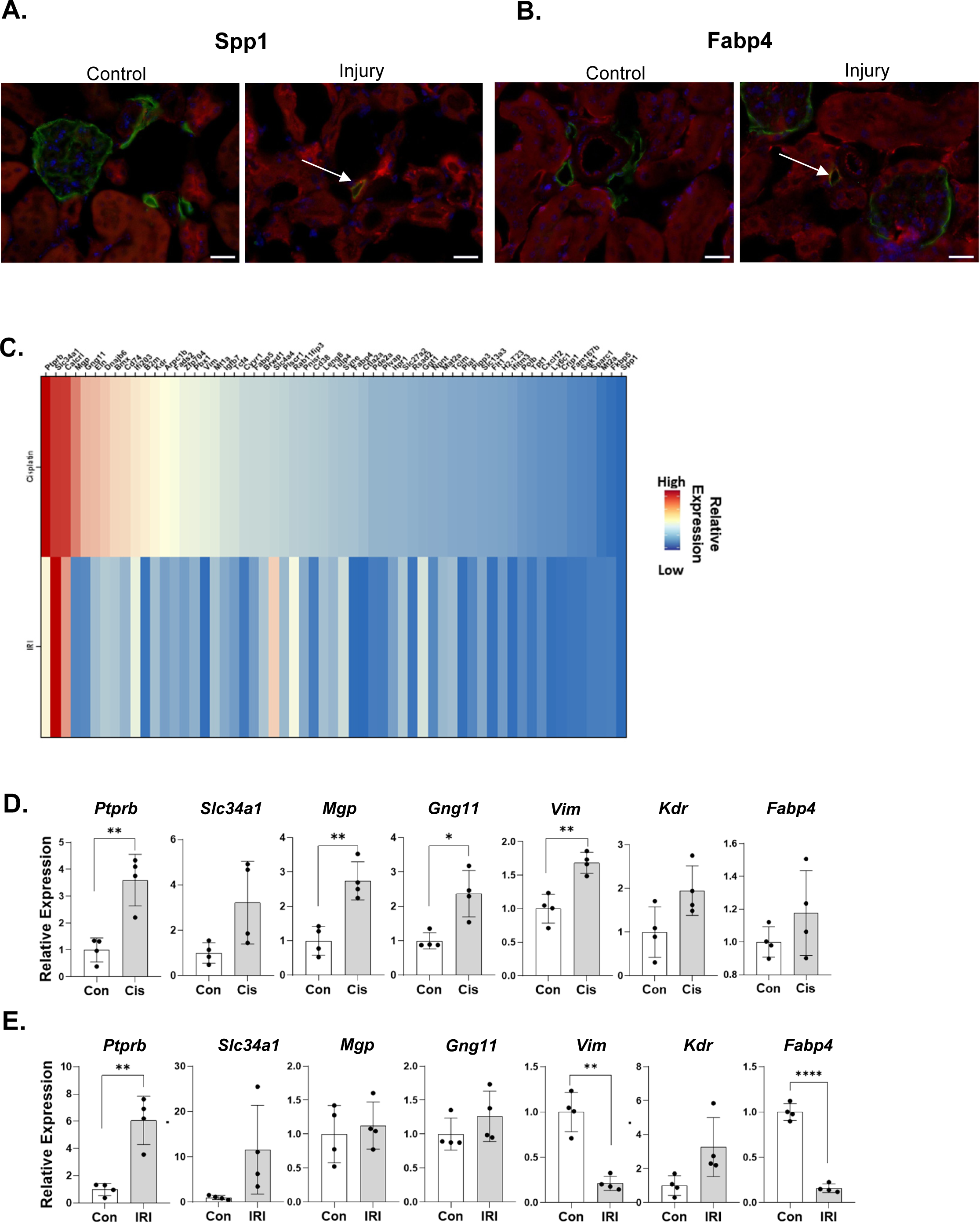
Validation of lymphatic genetic response to kidney injury. A) Immunofluorescent staining of Spp1 (red) and Pdpn (green) demonstrating increased Spp1 upon cisplatin injury and co-localization with Pdpn+ lymphatic vessel (arrow). B) Immunofluorescent staining of Fabp4 (red) and Ppdn (green) with co-localization in injury (arrow). Images collected at 40x on 10uM sections. Scale bar= 20uM. C) Heatmap of log2FC values of isolated renal LECs for highly changed genes as determined by scRNA sequencing data in cisplatin and ischemia reperfusion injury (IRI). Genes are normalized to respective condition controls. D) qPCR relative expression values in cisplatin isolated renal lymphatic endothelial cells (LECs) confirms significant gene expression changes with injury for *Ptprb, Mgp, Gng11,* and *Vim* as identified in scRNA sequencing results in*. Kdr* (p = 0.0575) and both *Slc34a1* and *Fabp4* were not detected as significantly different. D) qPCR relative expression values in renal LECs isolated from ischemia reperfusion injury (IRI) confirms similar significant gene expression changes with injury for *Ptprb* and similar trends to cisplatin LECs for *Slc34a1* and *Kdr.* In IRI *Fabp4* had significantly decreased relative expression when compared to control conditions. *= p <0.05, **= p <0.005, ****= p <0.0001.

To determine if detected gene expression genes were due to a direct cisplatin interaction with LECs, HDLECs were challenged with cisplatin *in vitro*. A heatmap of all tested genes normalized to saline controls indicated genes (*Ptprb, Vim, Kdr, and Fabp4)* were significantly increased with 8 µg/mL cisplatin (Figure 6A, B, Supplemental Table 6). However, *Slc34a1* and *Spp1* were significantly decreased with cisplatin treatment. These data indicate that cisplatin does not directly drive identified LEC genetic changes.

**Figure 6:**
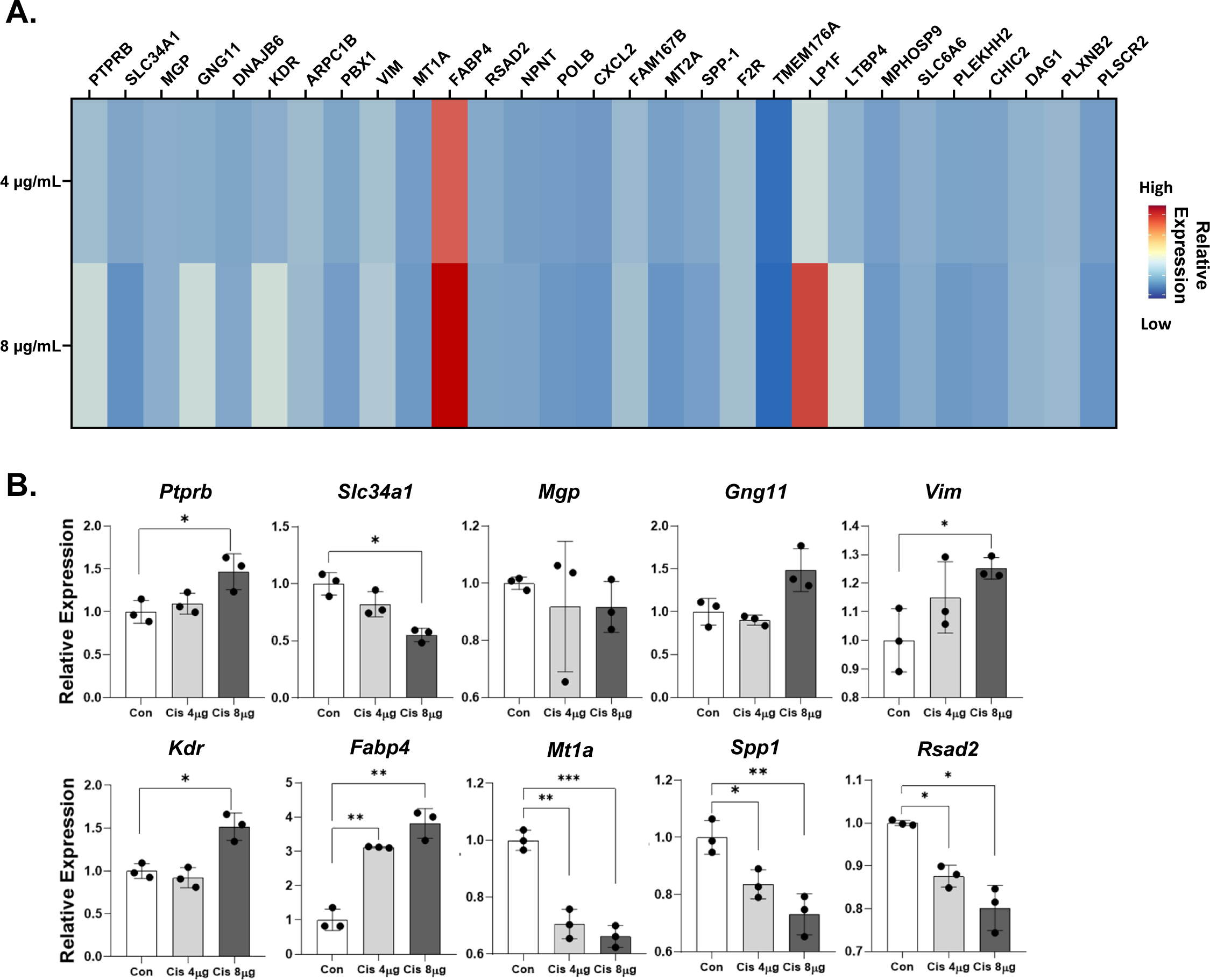
Gene expression of cisplatin-treated HDLECs *in vitro*. A) Heatmap of log2FC values of Human Dermal Lymphatic Endothelial Cells (HDLECs) highly changed genes as determined by scRNA sequencing data. HDLECs genes compared in cultures treated for 24 hours with 4 µg/mL or 8 µg/mL cisplatin. Genes are normalized to saline treated controls. B) Relative expression values of *Ptprb, Vim, Kdr, Fabp4* and *Rsad2* in HDLECs confirming scRNA sequencing gene expression. *Spp1* relative gene expression unlike scRNA sequencing expression and immunofluorescence validation. *= p <0.05, **= p <0.005, ***= p<0.0005.

## DISCUSSION

Several physiological and pathophysiological roles have been described for lymphatic vessels in the kidney, but renal LECs and their response to injury remains largely unstudied, in part due to their relative scarcity compared to other renal cells. Our study applied scRNA sequencing technology to characterize the renal LECs in quiescence and injury and has identified distinct LEC populations, potential molecular roles, and changes in genetic expression with renal injury, uncovering previously undefined roles for lymphatic vessels in the kidney injury response. We have also confirmed, as indicated in previous studies, that upon kidney injury tubules increase Pdpn expression^54^. Our data further confirms the LEC genetic response is likely injury context dependent, supported by our genetic confirmation in cisplatin and an IRI injury models.

In most tissues, lymphatic vessels begin as blind-ended capillaries that coalesce into pre-collector vessels, that possess unidirectional valves, and evolve further into larger conducting vessels. Several studies have characterized the heterogeneity of LECs along this network from the capillary tips to conducting vessels and highlight the transport and immunomodulatory roles of LECs^15, 24^. Past studies examining the anatomy of lymphatics in the kidney have identified a network of lymphatics along the interlobular and corticomedullary vasculature with some vessels noted in the murine cortex and subcapsular space; recent lightsheet microscopy of adult mouse kidneys beautifully demonstrated renal lymphatic anatomy^17–19^. Using gene expression from past LEC sequencing efforts, we were able to confirm populations of collecting vessel LECs, pre-collecting LECs, and LECs from lymphatic valves. Interestingly, indications of capillary LECs were few, but conform to the past anatomical descriptions of the network.

The growth of new lymphatic vessels, lymphangiogenesis, occurs post injury as an often-necessary step to resolve inflammation and restore tissue homeostasis ^55–57^. Several recent elegant studies have characterized lymphatics in the developing kidney^58 59^. The importance of renal lymphatic vessels in pathological progression in the adult kidney, however, and how manipulation of lymphangiogenic signaling through vascular endothelial growth factor receptor-3 (VEGFR-3) impacts functional outcomes, is less clear. What is clear is that the lymphatic vasculature and VEGFR-3 signaling is involved in transplant rejection, renal fibrotic remodeling, immune accumulation, and microvascular rarefaction^21, 60, 61^ ^62^. After AKI insult, expression of the lymphatic growth factors VEGF-C and VEGF-D, are increased within the kidney and most models (and clinical samples) demonstrate increased lymphangiogenesis at some point post injury, however the timing and extent of injury may likely regulate the importance of endogenous post-injury LEC responses and lymphatic impact^6, 9, 63, 64^. It is thus not surprising that a series of vasculogenic pathways were some of the most upregulated in LECs 3 days following injury in our data.

Lymphatic vessels are increasingly recognized as active regulators of the immune response^65–67^. For example, in cancer models LECs can upregulate PD-L1, present antigens through MHC expression, and impact tissue cytotoxic CD8+ T cell numbers homeostasis^66 68^. Studies in other disease models have demonstrated that LECs have a molecular repertoire which specifically alter immune cell activation, differentiation, trafficking, chemokine receptor expression, and antigen presentation^10, 66, 69^. Past work from our lab and others have demonstrated that manipulating renal lymphatic density alters immune cell populations post-injury^8, 9, 22, 62^. We therefore hypothesized that renal LECs alter their molecular roles in response to injury. Here we have confirmed renal LECs transcriptionally adapt to cisplatin and IRI, uncovering an underappreciated role of renal lymphatics as an active regulator of the response to injury stimuli. We find renal LECs upon injury stimuli have increased MHC gene expression (*B2m, H2-D1*),) and are involved in biological processes related to T cell differentiation (*Ctla2a, Id1, Cd38, Rsad2*), and antigen presentation and cytokine signaling (*Lyc61*, *Foxp1, Plscr2*). Whether renal LECs maintain the same molecular functions and immunoregulatory roles determined in this study across other kidney injury models remains an open question, but at least some of the repertoire was present in LEC-enriched cells from IRI kidneys. We propose that renal LECs are potentially regulators of inflammatory resolution post-kidney injury, and, importantly, that studies targeting LEC genes associated with immune cell interactions should be conducted to determine post-injury outcomes.

There are several potential limitations for the study. First, we utilized only male mice for renal LEC enrichment studies. It is likely gender plays a role in LEC specific responses as previous studies have identified female mice of C57Bl6/J background to have a degree of protection from AKI, likely an estrogen mediated effect^70, 71^. Second, due to the lack of renal LEC databases we identified and characterized clusters with the understanding they are CD31+Pdpn+ and utilized *Sox18, Prox1,* and *Flt4* as additional identification markers. To our knowledge there is currently no database or resource to specifically catalogue genes associated with renal LECs, although some studies have identified small LEC populations^12, 13^. Our study utilizes the LEC genetic signatures currently available, however, with the increasing resolution and capabilities of next generation sequencing, new lymphatic populations may be identified. Specifically, once de novo lymphatic vessels are formed post-injury (after day 7 according to other cisplatin studies) they would be presumably of the capillary type which have previously demonstrated a strong immunoregulatory pathway expression^15^. Finally, the cisplatin injury model introduces systemic as well as local effects which may contribute to the overall finding of the scRNA sequencing data set. We mitigate this shortcoming by assessing gene expression of isolated renal LECs in an IRI model and confirming cisplatin specific effects with HDLEC studies. Future studies would focus on applying scRNA sequencing in various injury models to determine if LECs alter response to injury dependent on the injury stimulus.

Lymphatic vessels and the LECs that make them up provide for an intriguing target to manipulate tissue pathological responses. The immunological mechanisms by which AKI progresses to CKD may be tied to LEC-immune interactions. The renal network of lymphatic vessels and LECs may provide a future target to change the inflammatory progression of AKI.

## Summary/Translational Statement

Lymphatic endothelial cells (LECs) in other disease models have been demonstrated to regulate the immune response and subsequent inflammation. In this study we utilized scRNA sequencing to investigate how LECs respond to renal injury. Our results indicate that renal LECs alter genetic expression associated with biological processes such as lymphangiogenesis and immunomodulatory interactions. Targeting LEC-immune signaling may provide a novel therapeutic target to limit post-AKI inflammation and progression to chronic kidney disease.

## ACKNOWLEDGEMENTS

The authors thank Andrea J. Reyna and countless undergraduates for animal husbandry support. Library generation and sequencing were performed at the Texas A&M Institute for Genome Sciences and Society (TIGSS) Experimental Genomics Core.

