## Supplemental Figure for "Single-cell RNA sequencing identifies response of renal lymphatic endothelial cells to acute kidney injury"

**List of Supplemental Tables**

**Supplemental Table 1: qPCR Primers**

**Supplemental Table 2: Antibodies Used**

**Supplemental Table 3: Control conditions global genes**

**Supplemental Table 4: Control LEC subcluster differentially expressed genes**

**Supplemental Table 5: Injury response differentially expressed genes**

**Supplemental Table 6: Control versus injury differentially expressed genes**

**Supplemental Table 7: qPCR data values with averages**

**Supplemental Legends and Figures**

**Supplemental Figure 1:** Identification markers for scRNA sequencing clusters. A) Gene markers used to identify vascular, hybrid vascular, and immune cell populations from unsupervised clustering analysis. B) Cluster assessment of previously published LEC genetic markers to distinguish capillary, pre-collector, collector, and valve LECs.

**Supplemental Figure 2:** Podoplanin labeling of other renal cells. A) Podoplanin labeling (green) across kidney regions also labels “hybrid vasculature” in the outer strip of the medulla (OSOM) with no other labeling in the cortex (C) or inner strip of the medulla (ISOM) to inner medulla region; In injury, podoplanin+ expression appear in the ISOM. Bars = 500 um. B) Podoplanin (green) and UTB/Slc14a1 (red) labeling of hybrid vessels in the ISOM in injury. C) Podoplanin (green) and PLVAP (red) labeling of hybrid vessels in the ISOM in injury. D) Podoplanin (green) and endomucin (red) labeling of hybrid vessels in the ISOM in injury. B-D Blue=DAPI. Bars= 200 um.

**Supplemental** **Figure 3:** LEC population global genetic response in quiescent and injury conditions. A) Volcano plots of significant log fold changes in LEC 1 between quiescent and injury injury illustrating significant gene changes compared globally B) Volcano plots of significant log fold changes in LEC2 between quiescent and injury conditions demonstrating globally significant gene changes. C) Global injury response heatmap of top 5 genes by scRNA sequencing identified clusters.

**Supplemental Figure 4:** Renal LEC GO biological pathways in injury. A) GO biological analyses of renal isolated renal LEC subset. B) MCODE interaction network of renal LEC genes indicating protein-protein vascular and lymphatic interactions. C) GO analyses of the Top 25 upregulated and downregulated injury LEC subset genes when compared to control LEC subset.

**Supplemental Figure 1**

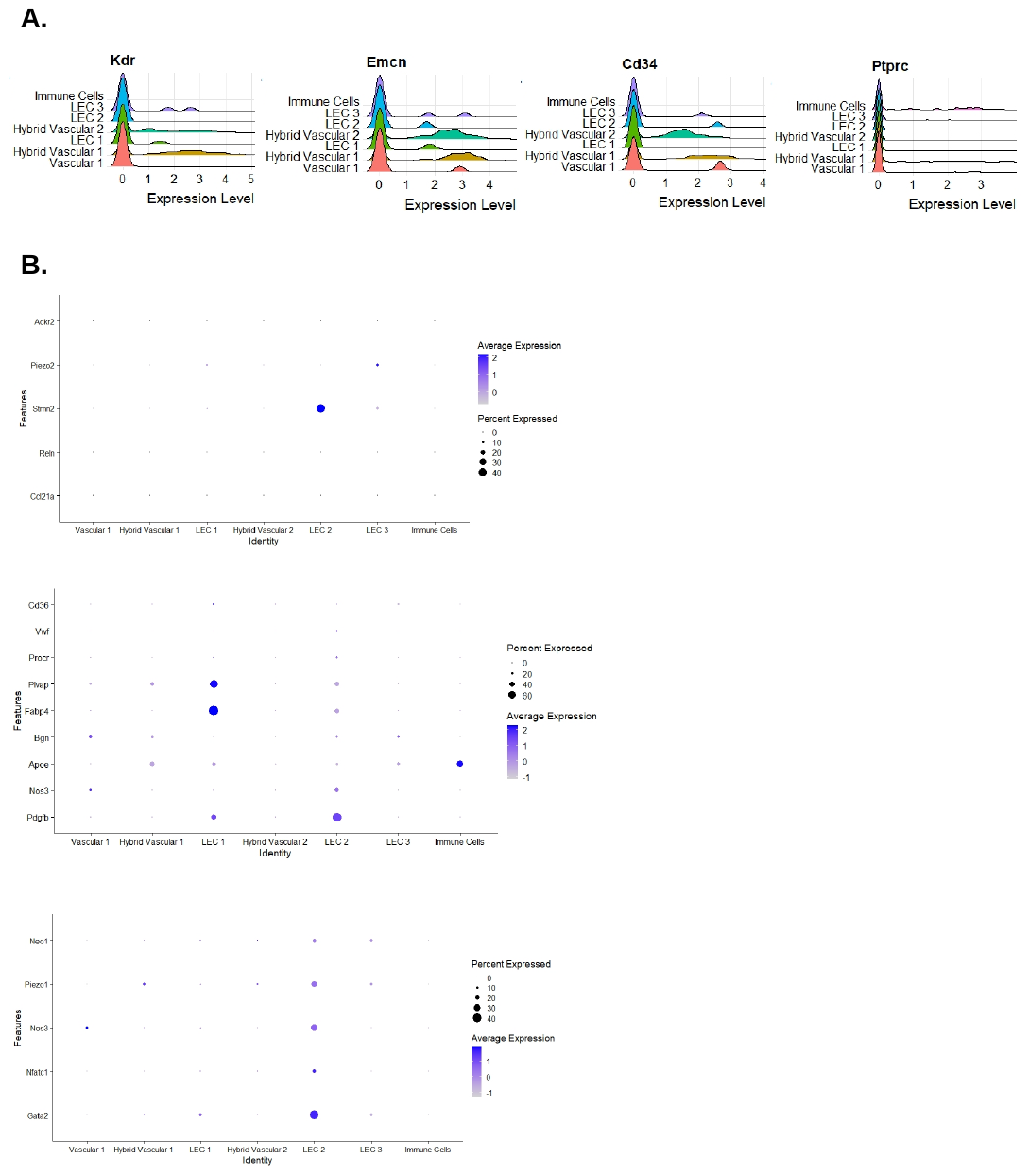

**Supplemental Figure 2**

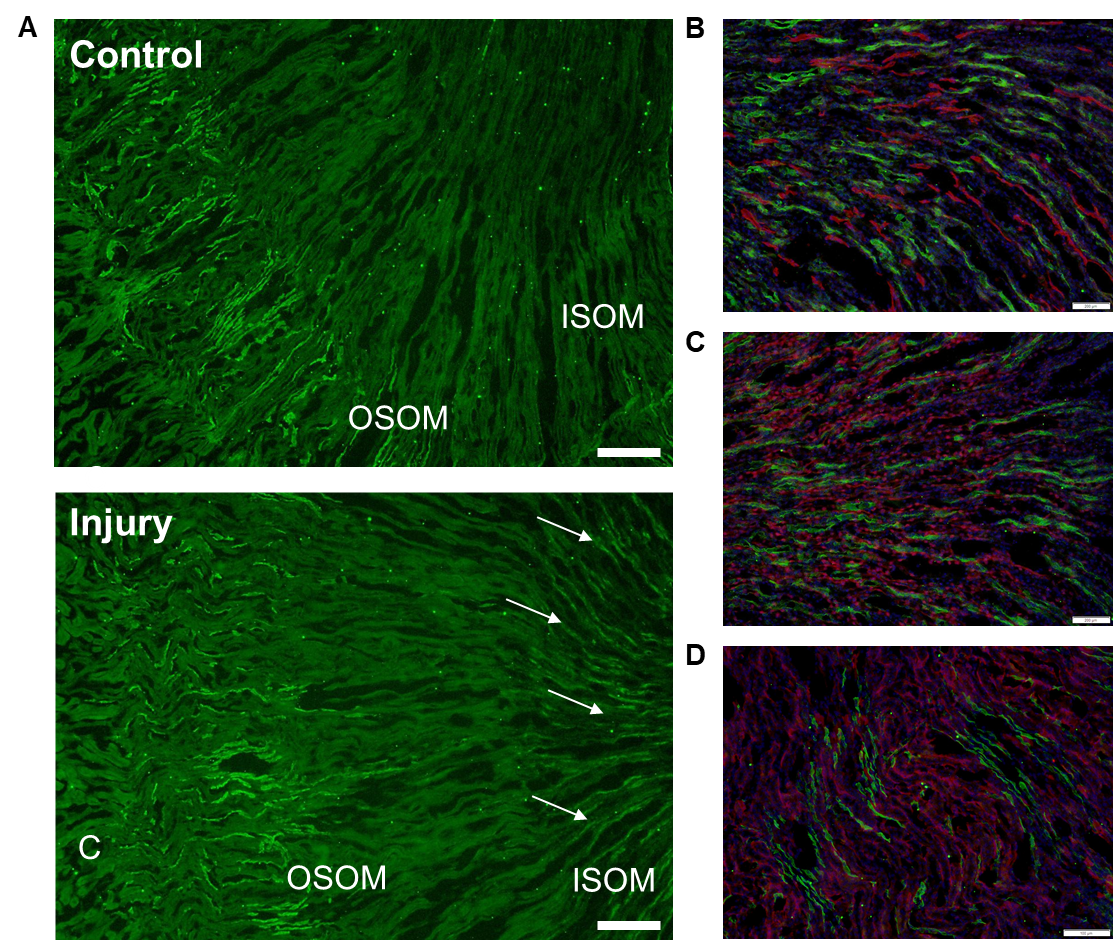

**Supplemental Figure 3**

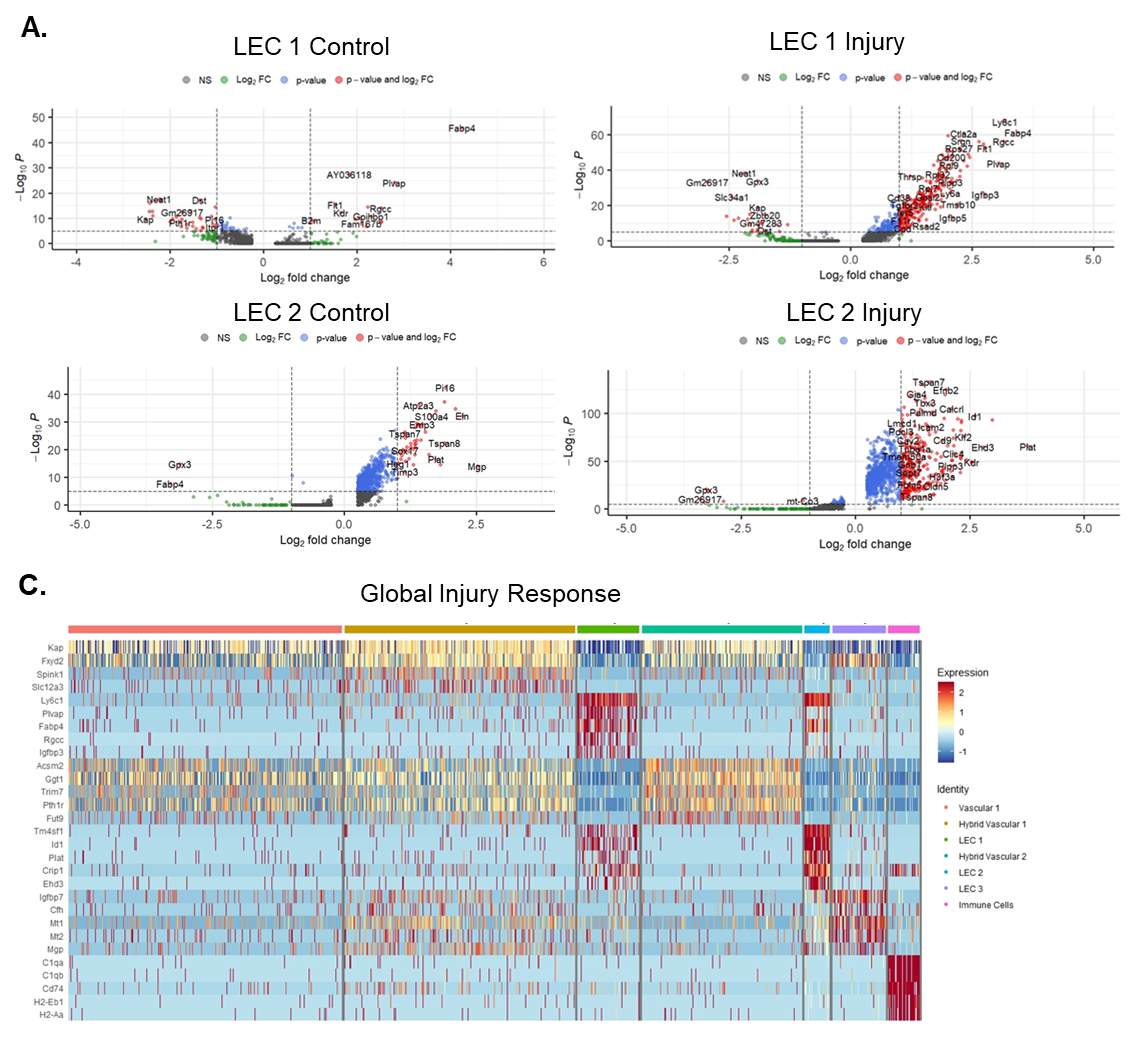

**Supplemental Figure 4**

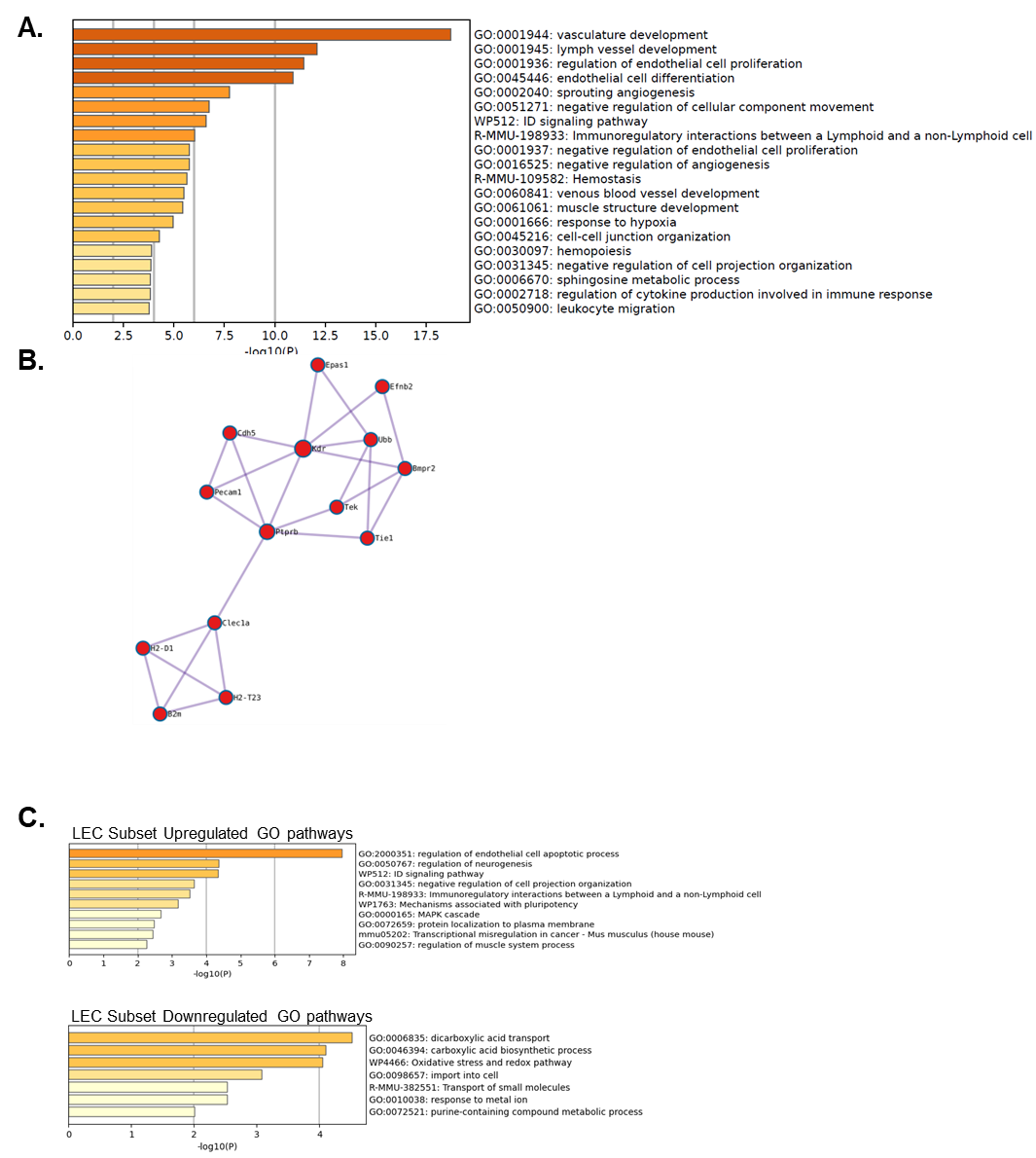
